# Imaging optically thick tissues simply and reproducibly: a practical guide to Lightsheet Macroscopy

**DOI:** 10.1101/2022.03.31.486601

**Authors:** Rebecca M. Williams, Jordana C. Bloom, Cara Robertus, Andrew K. Recknagel, David Putnam, John C. Schimenti, Warren R. Zipfel

**Affiliations:** BRC Imaging Facility, Institute for Biotechnology, Cornell University; Whitehead Institute for Biomedical Research; Meinig School of Biomedical Engineering, Cornell University; Meinig School of Biomedical Engineering, Smith School of Chemical and Biomolecular Engineering, Cornell University; Department of Biomedical Sciences, Cornell University

**Keywords:** Lightsheet Microscopy, Macroscopy, Tissue Clearing, Tissue Imaging

## Abstract

Lightsheet microscopy offers an ideal method for imaging of large (mm-cm scale) biological tissues rendered transparent via optical clearing protocols. However the diversity of clearing technologies and tissue types, and how these are adapted to the microscope can make tissue mounting glitchy and somewhat irreproducible. Tissue preparation for imaging can involve glues and or equilibration in a variety of expensive and/or proprietary formulations. Here we present practical advice for mounting and capping cleared tissues in optical cuvettes for macroscopic imaging, providing a standardized 3D cell that can be imaged routinely and relatively inexpensively. We show that acrylic cuvettes should be non-aberrating with objective numerical apertures less than 0.65, and present an inexpensive tool for alignment and calibration of standard lightsheet parameters. Mouse embryo, liver and heart imaging are demonstrated as examples with practical recommendations for acquisition and post-processing.

## 1. Introduction

Recent innovations in tissue clearing and optical imaging technologies have enabled optical macroscopy of large scale (mm - cm) pieces of fluorescent tissue, facilitating a transformative understanding of how proteins or cell populations are organized in 3D at the tissue and organ level.

The depth of imaging is primarily limited by optical scattering. A photon can only travel ∼100 um through tissue before it is scattered (mean free path, MFP). By ten times that distance (1mm), each photon has deviated from its initial path ∼ten times, so the information contained within the light is totally scrambled. Optical scattering generally limits the acquisition of clean, detailed images to a few hundred microns in biological tissues. Absolute numbers describing scattering coefficients (inversely related to the MFP) are extremely tissue and wavelength dependent. Reviews on tissue scattering [1-3] summarize these properties for a variety of tissues and wavelengths.

The refractive index (n) is a measure of how much the molecules in a medium interact with light. The imaginary part of this number quantifies absorption, whereas the real part quantifies a slowing of the light due to the aggregate oscillation of molecules and how well they track with the light wave frequency. This real part of the refractive index varies with the color of the light, and in biological tissues ranges from 1.0 in a perfect vacuum (no interaction) to 1.55 in hard tissues such as bone [4]. When light crosses an interface from one refractive index to another (such as water to lipid), part of it will be reflected and part of it will bend. When light encounters many of these interfaces in succession (optical scattering), it loses directionality and information, resulting in blurry, unfocused images.

Tissue clearing broadly encompasses a variety of protocols that reduce the amount and severity of these refractive index mismatches, leading to tissues that appear transparent. This collection of protocols enables miraculous imaging depths that are in some cases 100-fold longer that what would be possible in uncleared tissues (aka. whole mouse brains and organs, whole mouse embryos, whole plants). Tissue clearing is an old science; a method using Benzyl Benzoate and Methyl Salicylate was described as early as 1914 [5]. Research developments in the past decade have elucidated new clearing technologies that are now compatible with fluorescent tags and in some cases even fluorescent proteins. Though excellent reviews exist on the subject [6-8], the field is obfuscated by an exploding variety of acronyms, and in some cases proprietary formulations that are unknown and expensive.

The goal for this paper is not to simplify or advance any of the steps involved in tissue clearing, but to describe protocols that enhance reproducibility and ease-of-use, and decrease costs associated with lightsheet (LS) microscopy of cleared tissues, including mounting in disposable optical cuvettes.

## 2. Results

Lightsheet microscopy is an ideal method of imaging optically cleared tissues to centimeter-scale depths. This process might require, for example, thousands of images at 10 um intervals. A LS microscope is significantly faster than a more-standard raster-scanned confocal microscope (the full frame is illuminated at once), and it limits illumination (and thus photodamage) to the plane being imaged. So this technique is ideal for large cleared tissues, and relatively simple in terms of its required instrumentation.

### 2.1. The case for optical cuvettes

The practical difficulties with performing LS microscopy come almost exclusively with the mounting process. More standard microscopies only require optical access from a single side, which is usually coverslipped. LS microscopy instead requires optical access from at least two and sometimes three sides. The specimen has to be immobilized while maintaining this access and high-NA optics are not generally forgiving with respect to available working distances. Preparing samples within cuvettes can be useful for a variety of reasons. 1) The imaging chamber of a LS microscope may require over 100 ml of clearing solution to fill it, which can be so expensive that it is essentially infeasible. 2) Specimens within cuvettes can give a robustness and reproducibility to imaging protocols. Specifically, the initial equilibration of the imaging media with the specimen media can cause optical nonuniformities within the imaging chamber, with resulting imaging aberrations. Also, because biological tissues can be soft, fluidic motions due to focusing can cause specimen movement during imaging. Specimens within capped cuvettes can be prepared anywhere and easily transported and stored. Additionally, specimens can be imaged and re-imaged repeatedly with no tissue handling required. Disposable cuvettes are readily available and inexpensive.

For large cleared tissues, we recommend sample mounting in capped cuvettes with four clear sides. These can then be immersed in a less expensive imaging solution that is index matched to the clearing solution. In particular, 2,2’-thiodiethanol (TDE) is relatively non-toxic and inexpensive, and can be matched to 1.33 - 1.55 by dilutions in water *[9]*. Expensive clearing solutions in direct contact with the biological tissue are only required within the cuvette. We recommend “ UV grade” cuvettes because they are acrylic (Polymethyl methacrylate, PMMA) rather than polystyrene (PS) based, with an index of refraction that is closer to that of most tissues.

Figure 1 is an example showing a cleared and mounted mouse heart that is attached to our most utilized 3D printed stage adapter designed for integrating with the LaVision BioTec Ultramicroscope II. The surface .stl file for 3D printing this adapter can be accessed [here](https://cornell.app.box.com/s/0hp9882mvrslv6jbex8ex9glmyqt9tke).

**Figure 1.**
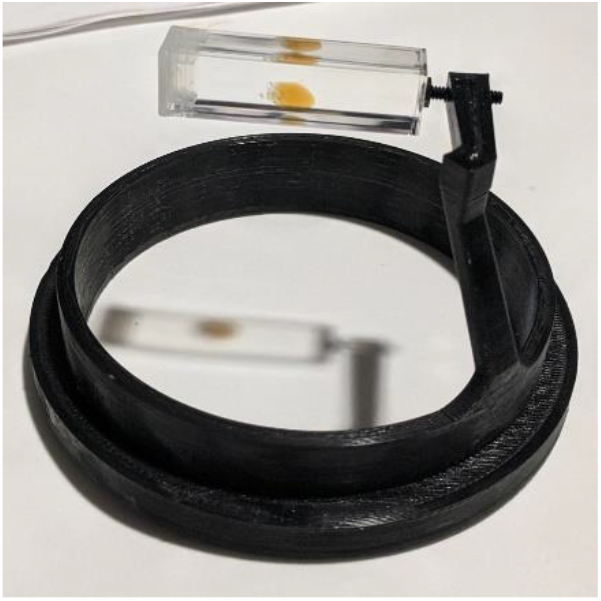
Cleared mouse heart in cuvette mounted on stage adapter.

### 2.2. Caveat with cuvettes: Spherical aberration

The primary optical aberration introduced by a flat window is spherical aberration due to an overall index mismatch caused by the window with respect to the surrounding optical fluid. Cuvettes with 3 or 4 clear sides are only available in 1 cm and 3 mm widths (each with 1 mm wall thickness leading to 1.2 cm and 5 mm total diameters). Lens working distances must accommodate the wall thickness as well as the sample thickness. Using cuvettes leads to a situation in which the refractive index of the outside imaging solution is matched with that of the tissue clearing solution, but the cuvette has an unmatched higher index. For example, our users frequently employ the DIX clearing solution developed by Murray and colleagues *[10]*, with an index matching solution of 60% TDE *[9]*, both with n=1.45. But disposable cuvettes are typically made of acrylic or polystyrene with refractive indices of n=1.50 and 1.55 respectively. This thickness of plastic (*t*) will cause a phase aberration given by *[11]*:

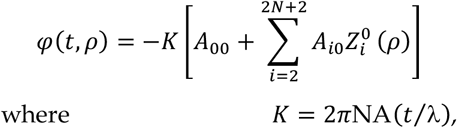

the 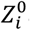 are Zernike circle polynomials of *i*^*th*^ order and zero kind (radially symmetric), and the *A*_*i*0_ coefficients are functions of the objective NA and the two unmatched indices of refraction *n* of the imaging solutions and *n*_*c*_ of the cuvette plastic:

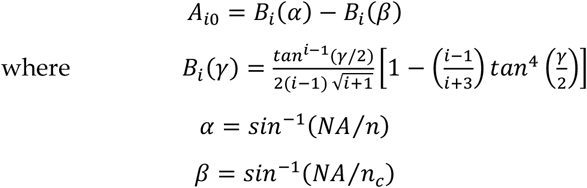

The point-spread-function (PSF) characterizing these aberrations can be calculated via standard methods [12]. The results are shown for varying amounts of first order spherical aberration (the 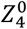 term, fig. 2A). The primary effect of spherical aberration is to lengthen the PSF axially, which expands the focal plane such that detailed information can no longer be acquired from a thin optical section. If the axial full-width-at-half-maximum (FWHM) is plotted as a function of the magnitude of this aberration, it can be seen that this lengthening of the PSF becomes significant at ∼*π*/4 (fig. 2B). As long as conditions are chosen such that *KA*_40_ is less than this value, the effects of spherical aberration are minimal. For real world examples, we assume a CLARITY-like mounting index (*n*=1.45, 450 nm to 1.46, 650 nm), a standard cuvette thickness of 1mm, and illumination wavelengths at two ends of the spectrum (450 nm, blue; 650 nm, red). The expected magnitude of spherical aberration is plotted as a function of NA for two standard cuvette plastics. Calculations demonstrate that it will be critical for researchers to choose acrylic based cuvettes for high-resolution imaging (NA>0.5), and this technique will not work for NA’s > 0.7. (Most objective lenses with NA’s this high would not have sufficient working distance for imaging through a cuvette wall anyway.)

**Figure 2.**
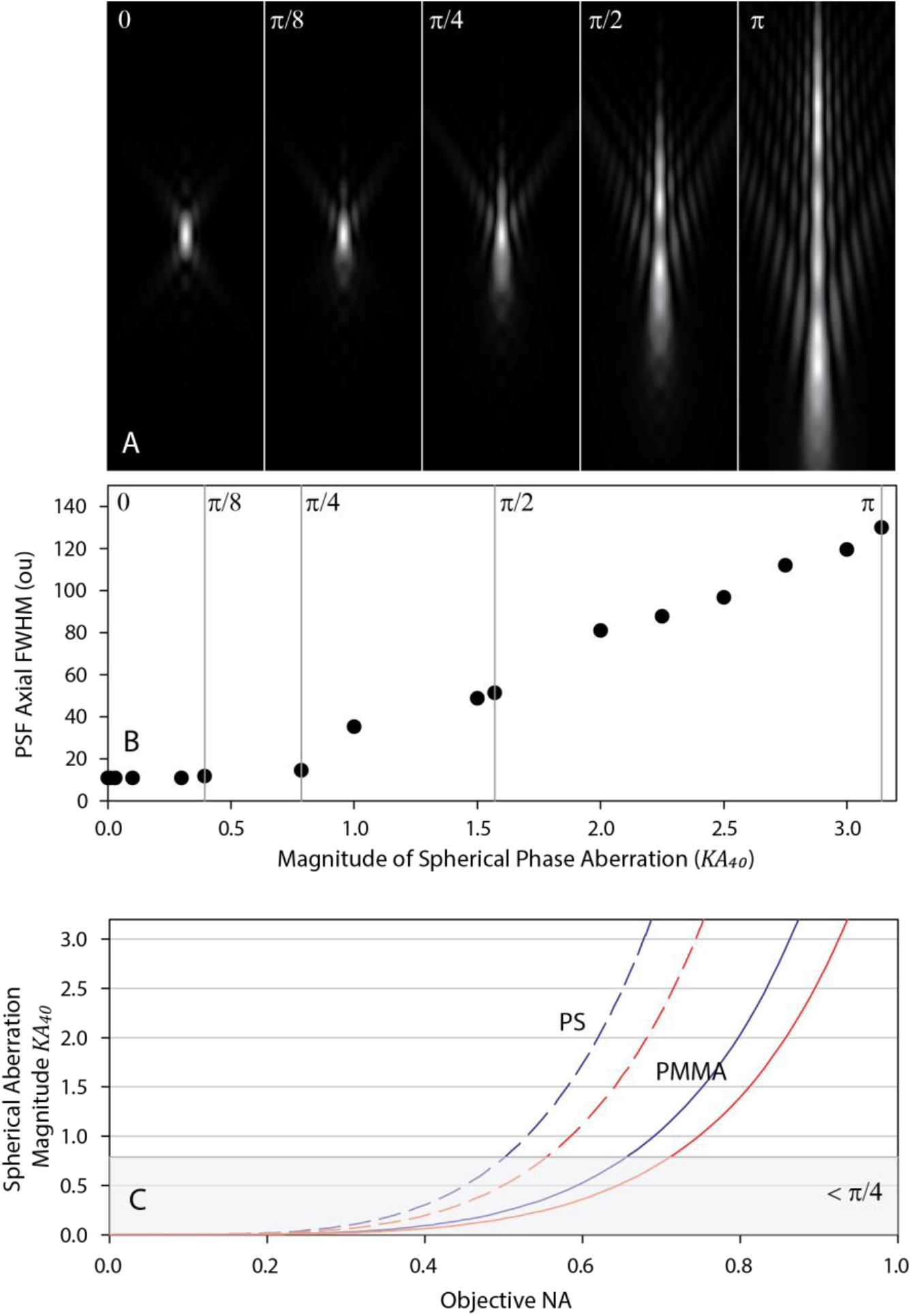
Effects of cuvette refractive index mismatch caused spherical aberration (SA) on lightsheet PSF. A) Axial cross-sections of PSF’s with indicated amounts of SA. B) The axial FWHM is relatively constant up to a magnitude of π/4 SA. C) Calculated SA magnitude from index mismatches due to cuvette walls (1 mm material) of two standard cuvette plastics: PMMA (solid lines) and PS (dashed lines) at blue and red wavelengths. SA remains below acceptable levels for NA<0.5 (PS cuvettes) and NA<0.65 (PMMA cuvettes).

### 2.3. Finding the lightsheet focus

With a standard microscope, the user focuses on the sample. With a LS microscope, the user visually assesses whether the LS is in the sample, and then focuses to the LS using the viewing objective. Once this is done, there is no more focusing with the viewing objective, only moving the sample relative to the LS and imaging objective. Adjusting the LS focus to the field-of-view (usually the X direction on most setups) needs to be adjusted for each sample type, and can often be difficult to discern qualitatively. Ideally in a LS microscope, there would be an infinitesimally thin sheet that extends infinitely. This is the reason that some more complex LS microscopes employ Bessel beams *[13]*, which are non-diffractive but carry extra fringes whose effects must be mitigated via other means. Using standard optics, a tightly focused beam axially (Z, perpendicular to the imaging plane) means a tightly focused beam laterally (in X). As a result, a thin illumination sheet (high NA sheet optics) comes at the expense of a limited field-of-view (FOV). Fig. 3 shows an example of imaging with a tight illumination sheet (NA=0.153) that is only focused in the center of the FOV. A technical difficulty often encountered is that the location of the LS focus can be difficult to discern, especially in samples with discrete labeling, such as the fluorescent beads shown. At any imaged plane in this stack, beads appear circular, with similar sizes (fig. 3A). The intensity does change outside the LS focus, but intensity changes may not be evident in a real sample with puncta of different brightnesses. As expected outside of the LS focus, the axial extent of the beads is significantly elongated. Having a focused lightsheet is thus critical to the detail that can be extracted in a particular location. A significant axial blurring will be evident outside of the sheet focus. The positioning and size of the LS focus cannot be identified reproducibly with the 2D image only.

**Figure 3.**
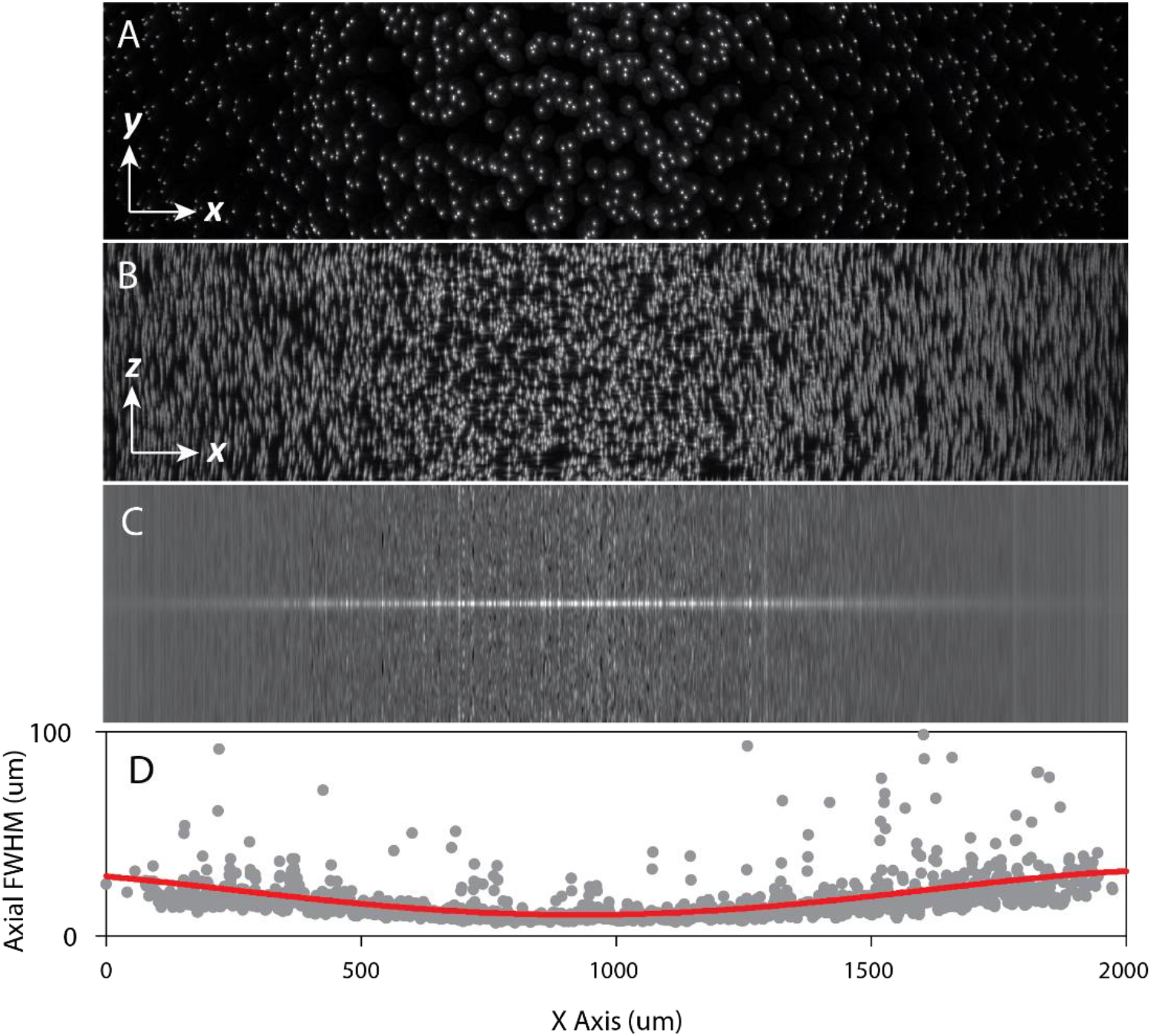
Lightsheet NA set such that the sheet is only focused in the center of the field-of-view (FOV) of gel-embedded beads. The dataset consists of a 3D volume (X: 2160 um x Y: 2560 um x Z: 500 um) in which the zoom and Z-step are chosen so that the voxel size is isotropic with 1 um spacing. A) XY maximum intensity projection through Z (only the central 500 um in Y is displayed). B) XZ maximum intensity projection through Y of 2 um fluorescent beads embedded in an agarose gel. C) Image in (B) autocorrelated in Z (each vertical column), demonstrating the positioning of the focused light sheet. D) A Gaussian fit to each column is used to determine the average axial FWHM of the 3D PSF’s. Decreasing the NA would lengthen the focus (∼1/NA^2), at the expense of slightly thickening the sheet (∼1/NA).

As displayed in figure 3, understanding and setting LS parameters can be done by 3D imaging with a bead impregnated gel. In this case the 3D PSF’s can be determined throughout the entire 3D field, each PSF resulting from a convolution of the excitation sheet intensity with the PSF of the imaging objective. However this protocol requires significant expertise and effort for data acquisition and analysis. These type of metrics are too complex to be performed for setting day-to-day optimization.

### 2.4. Simple Alignment Tool

Alignment parameters of the LS microscope can be somewhat difficult to set routinely and reproducibly. Alignment tools can be expensive and perhaps more importantly are often constructed with imaging solutions different from the clearing solutions used by individual researchers. Here we describe an inexpensive and simple calibration and alignment tool that is useful for adjusting day-to-day parameters on the LS. The calibration tool consists of a cuvette with a plastic slide cut to fit diagonally within the cuvette, painted with a commercially available fluorescent marker and filled with the clearing solution of choice (fig. 4A).

**Figure 4.**
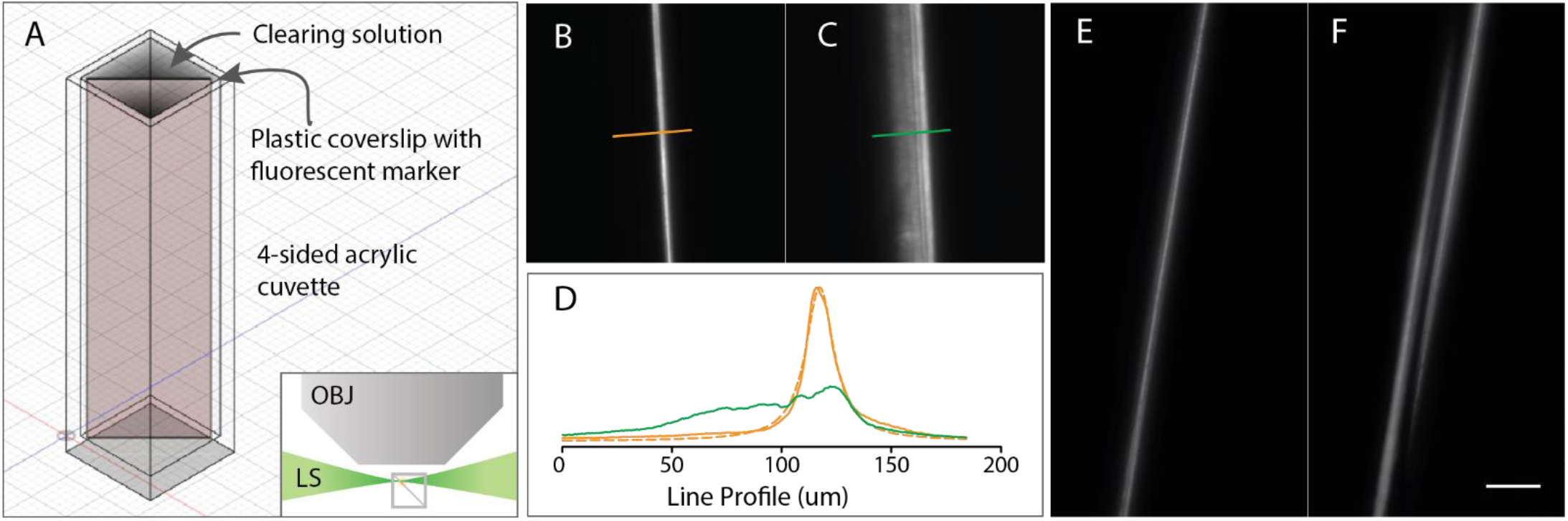
Simple lightsheet calibration tool constructed with a plastic cuvette and a plastic coverslip, painted with a fluorescent marker on one side. A) Diagram of tool and orientation (LS: lightsheet, OBJ: objective). B,C) Images acquired of the tool with the lightsheet focus adjusted correctly and incorrectly, respectively. D) Line profiles from images in (B, orange) and (C, green). The orange profile shows a Lorentzian fit with a width of 7.8 um (dashed line). E,F) Images acquired with left and right light sheets aligned and unaligned, respectively. The scale bar is 100 um.

The advantage of this tool is that it translates Z-information with respect to the LS thickness into XY information that can be read out immediately via the image. As previously demonstrated, the LS focus is difficult to visualize in samples with punctate fluorescence information because imaged points do not expand in XY with a thicker light sheet in Z. Figures 4B,C show images of this device when the horizontal focus is adjusted correctly and incorrectly, respectively. In the first case, the line profile of the coverslip surface can be well fit by a clean Lorentzian function (orange: data:solid and dashed:fit), whereas an out-of-focus LS significantly extends the thickness of the coverslip that is excited (green, fig. 4D). This tool is also extremely useful for aligning lightsheets that are originating from different directions. Figure 4E shows a blended image with two (+X and –X) lightsheets aligned, and Fig. 4F shows an equivalent image with the two lightsheets separated by 50 um in Z.

### 2.5. Autofluorescence

Autofluorescence is either a tremendous impediment or a tremendous asset in tissue imaging, depending on the user’s point of view. It can be used to form non-specific tissue images that can be extremely useful for providing the tissue context in which fluorescence is located. Because tissue autofluorescence is typically characterized by an emission spectrum that is broader than that exhibited by specific fluorophores [14], users can simply choose unused spectral bands for imaging autofluorscence, even at relatively long wavelengths (given sufficient power). For example, the location of GFP expression might not be obvious in the GFP channel alone (fig 5A). However imaging an “ off-GFP” channel (fig. 5B) shows only autofluorescence, such that the GFP signal can easily be identified in a merged channel (fig. 5C). Surplus autofluorescence in the gut can be decreased by feeding mice chlorophyll-free chow [15], or generally by a an overall decolorization (bleaching) of the tissues [6]. (Clearly this step cannot be used with endogenous fluorophores, because it does not distinguish autofluorescence from fluorescence.) Our strategy has been to utilize this signal by choosing excitation and emission filters that are distinct from the highly specific wavelengths associated with tagging fluorescent proteins or dyes. For example we used a 640nm excitation beam for generating the autofluorescence image in fig 5B. When merged (GFP:green; autofluorescence:violet), specific GFP expression becomes identifiable. Note imaging artifacts associated with a reduced transmission of shorter vs longer excitation wavelengths (green autofluorescence more evident at surfaces) to be discussed in the next section.

**Figure 5.**
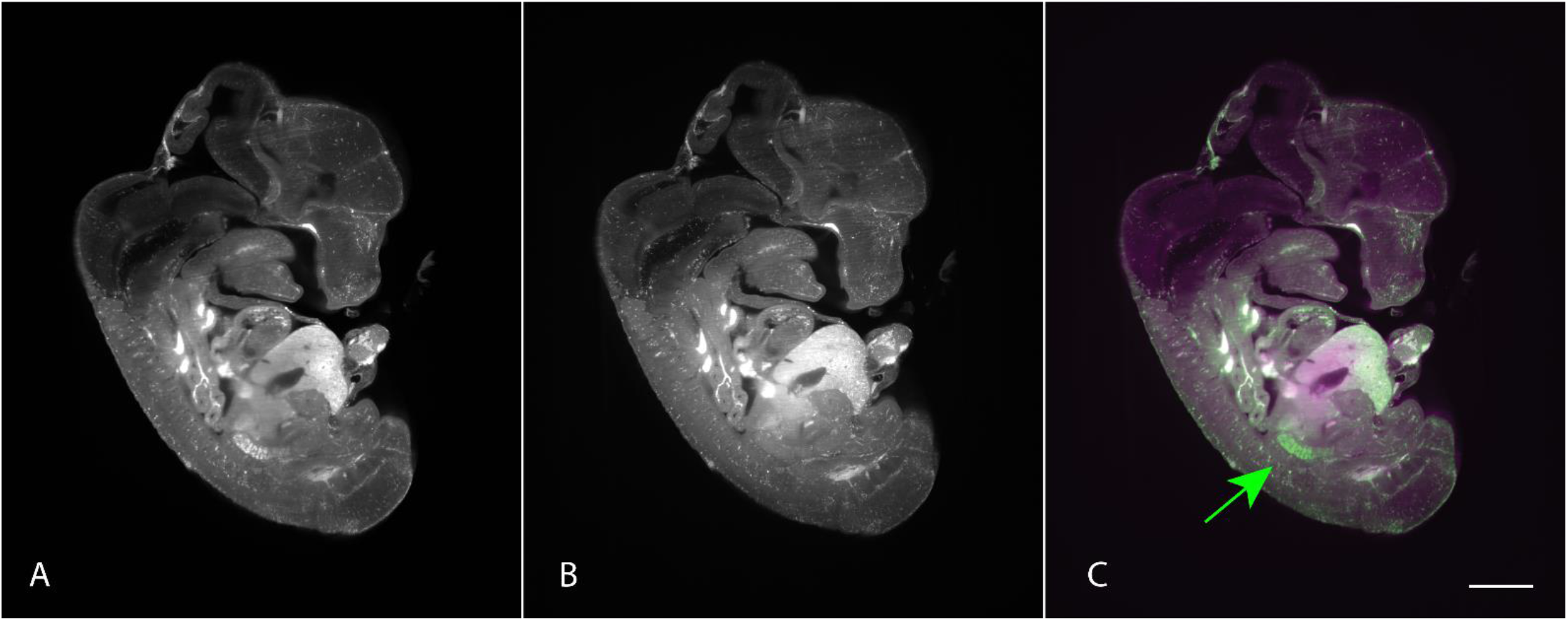
Sagittal image from a cleared mouse embryo (e12.5) expressing GFP in developing testes tissues. A) GFP channel (488nm excitation with a 525/50 nm bandpass) GFP expression is not at all obvious when only imaging in this channel. B) Autofluorescence channel (639 nm excitation with a 680/30 nm bandpass). C) Merging the two channels (GFP: green and autofluorescence: violet) enables identification of specific GFP expression (green arrow) within the overall tissue context. The scale bar is 1 mm.

### 2.6. Advantages of longer emission wavelengths

Scattering lengths are dependent not only on the quality of the cleared preparation, but largely on the excitation and emission wavelengths, with penetration depths typically increasing threefold from blue (488nm) to red (630nm) wavelengths [1]. These effects are clearly accentuated by high NA, simply because higher angles imply longer traveling distances within the tissue. Users must be wary of edge artifacts due to incomplete penetration of different colors to and from the interior of the tissue. This effect is evident in the gut in fig. 5, in which green signal is pronounced at the surface as compared to the red signal. Thus given the choice, users should choose fluorescent indicators with wavelengths that are as long as possible. In liver tissue (fig. 6), the structure of the parenchyma can be resolved at the surface of the organ, but very little detail is visible inside (680/30 nm emission, fig. 6B). However near-IR-labeled copolymers of quinidine and acrylic acid (845/55 nm emission) that preferentially accumulate in the liver (fig. 6C) are well resolved even within its depth.

**Figure 6.**
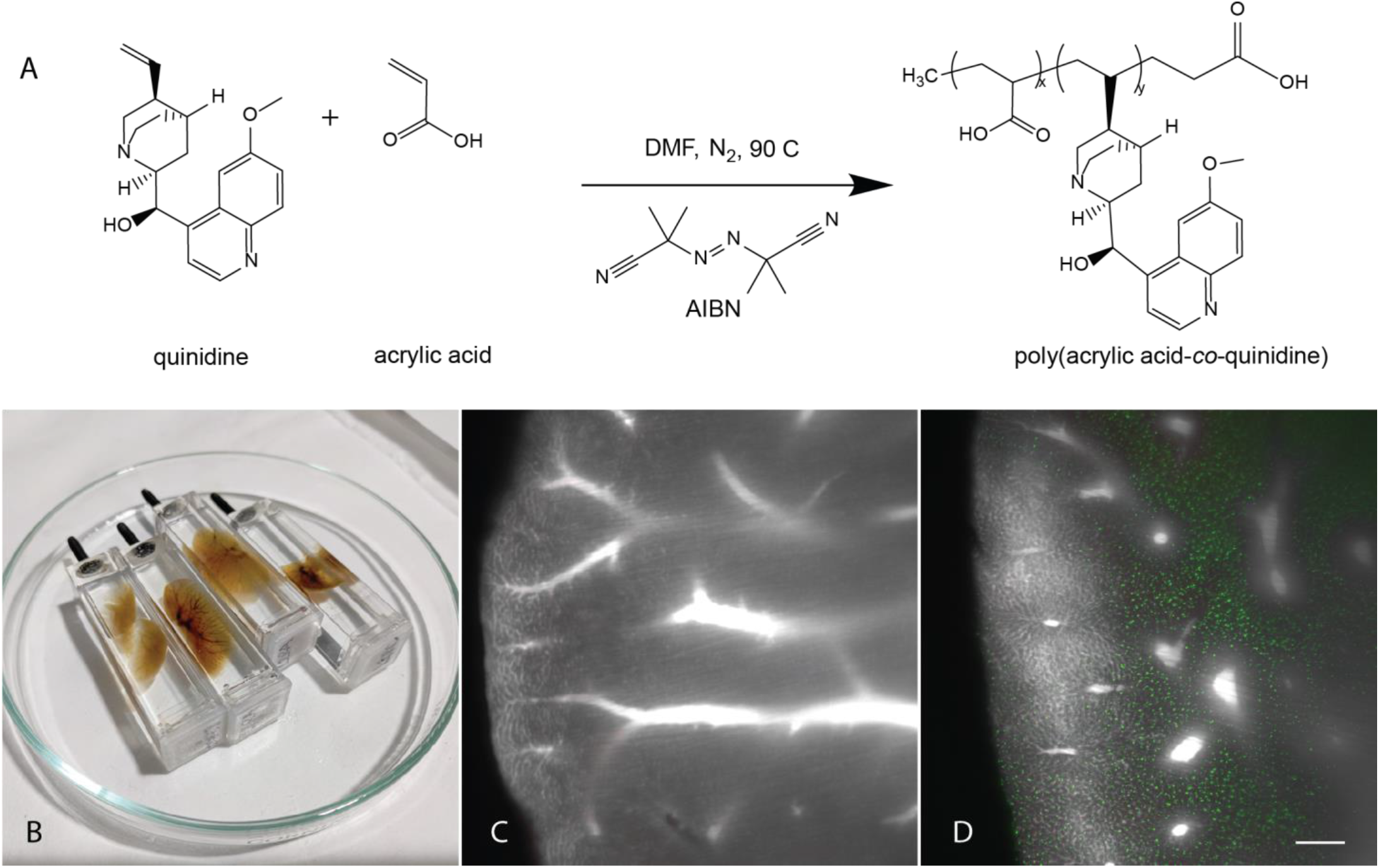
Cleared mouse liver samples exhibiting autofluorescence (greyscale) and Cy-7-labeled quinidine-acrylic acid copolymers (pseudocolored green). A) Reaction scheme for the production of a quinidine-acrylic acid copolymer via free radical copolymerization. B) Photograph of cleared mouse livers (right lobe and control snips). C,D) 40 um projections through a 3D ligthsheet stack from livers in which poly(acrylic acid) (negative control) and quinidine-acrylic acid copolymer were injected into the mice, respectively. The quinidine-acrylic acid copolymer preferentially accumulates in the liver parenchyma. Because these polymers were engineered to emit at extremely long wavelengths (845/55 nm emission), polymers are imaged well even through to the center of the organ. The scale bar is 100 um.

### 2.7. Mitigating streaking artifacts

LS images often exhibit classic streaking artifacts. These originate either from fixed scattering or absorption centers in the specimen or elsewhere along the optical excitation pathway, such as bubbles or refractive index changes in the imaging fluid. These nonuniformities distort or absorb the excitation beam before its focus, resulting in streaks corresponding to the geometric pathway of the excitation beam. Streak reduction is accomplished by either dithering the beam or constructing a sheet from several directions. In the Ultramicroscope II used for the imaging in Fig. 7, the focus is the aggregate of three beams that merge at the focus. Streaks such as these can be detrimental to clean segmentation and analysis of 3D objects within the dataset. Selected removal of discrete Fourier components associated with these streaks can in certain cases greatly aid the efficacy of downstream analysis. A ImageJ macro for removing these components through a Tiff stack of images is available [here](https://cornell.app.box.com/s/o2weqcm9y0af1kuvn0g9mnd6fowmf9y9).

**Figure 7.**
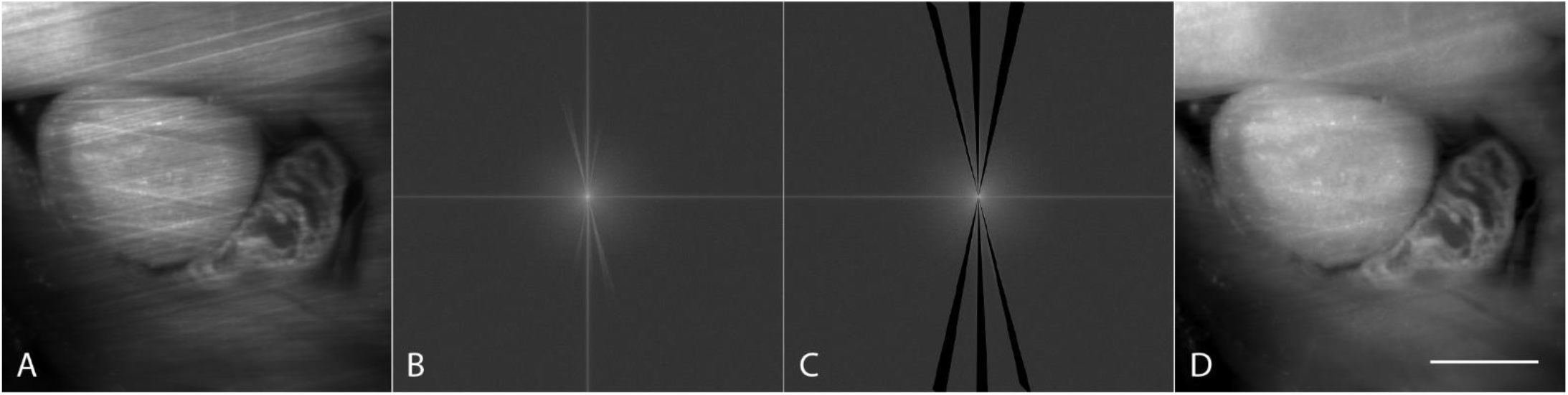
Removal of specific Fourier frequencies for mitigating LS streak artifacts. A) Cropped image from a cleared e12.5 mouse embryo detailing the developing heart (autofluorescence channel only) and typical streak artifacts. B) Fourier transform of (A). C) Removal of frequencies corresponding to geometric pathway of three converging lightsheets providing illumination for the image. D) Image resulting from the inverse transform with streak artifacts mitigated. The scale bar is 1 mm.

## 3. Discussion

Lightsheet microscopy offers an ideal technology for 3D imaging of optically-cleared, fluorescently-labeled tissues because it is fast (in most cases not limited by laser scanning) and the geometry of the sheet only illuminates the sample in the imaging plane, minimizing photobleaching. The LS geometry is also advantageous because optical resolution is inherently worse along the optic axis as compared to other axes (∼*λ*/NA laterally vs ∼*λ*/NA^2^ axially). Because the excitation and emission axes are perpendicular to each other, resolution can be more isotropic, leading to cleaner datasets with a less elliptical PSF, especially with low NA optics.

Practical difficulties with using this technique often stem from mounting tissues while maintaining optical accessibility along at least two optical axes, requiring protocols and expertise that depend specifically on the multitude of clearing protocols in use. Within this paper we described methods for water-based protocols using readily available and inexpensive optical cuvettes. With samples mounted in cuvettes, imaging protocols and procedures can be standardized, leading to enhanced reproducibility and ease-of-use. No part of the organ needs to be given up for attachment or stabilization glues or because of instability while imaging. We have shown that cuvette-based imaging is applicable to macro imaging with NA’s <0.5-0.6, depending on particulars associated with the clearing refractive index and cuvette material. For standard refractive indices ∼1.45, we recommend using acrylic-based cuvettes with lower refractive indices that better match standard clearing solutions (n∼1.45). We have described an inexpensive calibration standard for easily measuring the focus of the LS and aligning several lightsheets if they are available. These LS characteristics are critical to the quality of a 3D dataset and often difficult to determine in real samples.

Because autofluorescence tends to exhibit wide spectral characteristics, it can often be accessed in unused fluorescent channels. Spectrally broad autofluorescence can be invaluable for providing the tissue context for where the narrower specific indicators are emitting. Distinguishing fluorescence from autofluorescence requires logical controls. Fluorescence is usually in a single channel, whereas autofluorescence spreads between all channels. Related is the user’s ability to identify surface artifacts, especially evident in the shorter wavelength channels in which penetration to the center of the specimen is less prevalent due to higher scattering at these wavelengths.

Artifactual image streaking can ruin 3D dataset quantification, making accurate object segmentation difficult. Subtracting out Fourier components associated with streaking artifacts can smooth out images and enable a cleaner extraction of segmented objects.

## 4. Materials and Methods

Lightsheet images were acquired using a ***LaVision BioTec Ultramicroscope II***, using the MVPLAPO 2x objective on the Olympus MVX-10 zoom body with the corrected dipping cap. Images were acquired with an Andor Neo sCMOS camera. In this system, LS artifacts such as streaks and shadows are ameliorated by the use of six convergent light sheets (up to two triple light sheets from either side). The surface file for the 3D printed adapter for cuvettes can be accessed [here](https://cornell.app.box.com/s/0hp9882mvrslv6jbex8ex9glmyqt9tke). The printing fill factor should be as high as possible for arm stiffness as well as for providing a secure substrate for tapping the screw.

***3D point-spread functions (PSFs)*** were calculated using custom software written using the IDL image analysis environment (Harris Geospatial). PSFs were measured using 2.5 um green beads embedded in an agarose gel at 3.2 zoom and a 0.153sheet NA. The beads (488/515 LinearFlow cytometry beads, L14821 Invitrogen) were diluted 1/200 in 2% agarose and imaged in PBS with 488nm excitation and a 525nm (50nm width) emission filter.

A ***simple lightsheet calibration tool*** was prepared using a PMMA UV fluorometer cuvette (Perfector Scientific) with a plastic coverslip (Fisher Scientific 12-547) marked with an Expo Neon dry erase marker. The cuvette was then filled with refractive index matching solution in order to mimic standard tissue samples imaged in the facility.

All ***tissue imaging*** was done using clearing protocols previously described. In brief, lipids were removed from paraformaldehyde fixed tissues using 6% SDS (Sigma L3771) in PBS over a period of days to weeks, the length of time determined by a visual assessment of “ clearing”. The tissues were then rinsed for several hours in PBS and immersed in DIX refractive index matching solution [See table 1, 10] to equilibrate for one day.

**Table 1.** DIX index matching solution (per ml water).

| Compound | Sigma# | Concentration |
| --- | --- | --- |
| 60% iodixanol | D1556 | 0.30 ml/ml<br>(18% iodixanol) |
| n-methyl-d-glucamine | M2004 | 0.2 g/ml |
| diatrizoic acid | D9268 | 0.25 g/ml |
| sodium azide | S2002 | 2 mg/ml |

For mounting, 1.5% agarose was dissolved into DIX and heated, mixed and sonicated (Branson 1210 bath sonicator) until no particle scattering was observed. This solution was introduced into the cuvette with the sample to be imaged, and capped carefully such that no air bubbles were introduced. The orientation of the sample could be adjusted simply by tipping the cuvette until areas of interest were located near the cuvette wall, and the cuvette was then allowed to gel at room temperature. Methacrylate UV fluorometer cuvettes (Perfector Scientific 9014) were used in all imaging studies. In order to match the refractive index of the DIX (n=1.45), TDE (2,2′-Thiodiethanol, Sigma 88561) was diluted 60% in PBS and used in the imaging chamber outside the cuvette. This imaging solution was put aside, filtered and reused for multiple imaging sessions.

### Mouse embryos

Transgenic mouse strain B6;CBA-Tg(Pou5f1-EGFP)2Mnn/J, commonly referred to as Oct4ΔPE-GFP, (Jackson Laboratory 004654) was used in the embryo imaging experiments. Embryo dissections were performed 12 days after observing the presence of a copulatory plug.

### Mouse livers

Six-week-old female nude mice (Jackson Laboratory 007850) were used in the polymer biodistribution experiments. Polymers were administered via tail vein injection, and organs of interest were harvested after 24 hours. Synthetic methods for production of quinidine-acrylic acid copolymers are available upon request, and a reaction scheme is shown in Fig. 6.

## 5. Contributions and Funding

### Author Contributions

R.W. was responsible for conceptualization, methodology, data acquisition, visualization and analysis. J.B., J.S., C.R., and D.P. provided biological specimens and prepared specimens for imaging. A.R. assisted in the clearing protocols and imaging. W.Z. assisted in data analysis. All authors have read and agreed to the published version of the manuscript.

## Acknowledgements

Imaging was carried out with resources and expertise in the Cornell Biotechnology Resource Center (BRC) Imaging Facility.

## Institutional Review Board Statement

The use of mice in this study was approved by Cornell’s Institutional Animal Care and Use Committee (protocol #’s 2004-0038, 2011-0006 and 2018-0088).

